# Environmental transmission of *Pseudogymnoascus destructans* to hibernating little brown bats

**DOI:** 10.1101/2021.07.01.450774

**Authors:** Alan C. Hicks, Scott Darling, Joel Flewelling, Ryan von Linden, Carol U. Meteyer, Dave Redell, J. Paul White, Jennifer Redell, Ryan Smith, David Blehert, Noelle Rayman, Joseph R. Hoyt, Joseph C. Okoniewski, Kate E. Langwig

## Abstract

Pathogens with persistent environmental stages can have devastating effects on wildlife communities. White-nose syndrome (WNS), caused by the fungus *Pseudogymnoascus destructans*, has caused widespread declines in bat populations of North America. In 2009, during the early stages of the WNS investigation and before molecular techniques had been developed to readily detect *P. destructans* in environmental samples, we initiated this study to assess whether *P. destructans* can persist in the hibernaculum environment in the absence of its conclusive bat host and cause infections in naive bats. We transferred little brown bats (*Myotis lucifugus*) from an unaffected winter colony in northwest Wisconsin to two *P. destructans* contaminated hibernacula in Vermont where native bats had been excluded. Infection with *P. destructans* was apparent on some bats within 8 weeks following the introduction of unexposed bats to these environments, and mortality from WNS was confirmed by histopathology at both sites 14 weeks following introduction. These results indicate that environmental exposure to *P. destructans* is sufficient to cause the infection and mortality associated with WNS in naive bats, which increases the probability of winter colony extirpation and complicates conservation efforts.

## INTRODUCTION

Pathogens with indirect transmission from environmental reservoirs can have serious consequences for wildlife host populations (1). Environmental reservoirs can maintain infection in the absence of focal hosts, linking otherwise disconnected individuals across space and time (2-6). Furthermore, environmental reservoirs can sustain seasonal outbreaks (7-9) and increase the magnitude of disease impacts (10). For numerous diseases, including Chytriodiomycosis in amphibians (11), anthrax in ungulates (12), and white-nose syndrome in bats (13), population recovery may be limited by the continued exposure to environmental pathogen reservoirs.

White-nose syndrome (WNS) is a disease of hibernating bats first documented in 2006 in eastern New York State, USA (14). It has since spread across much of North America (13) and threatens multiple bat species with extinction (15). In New York and Vermont, the states with the longest history of WNS, the numbers of bats in hibernacula have declined overall by more than 95% (13, 15). White-nose syndrome is caused by the psychrophilic fungus *P. destructans* (16), which appears to have been introduced to North America from Eurasia (17). This fungus invades living tissue of torpid bats (18) and disrupts the normal pattern of periodic arousal in hibernating bats (19). *Pseudogymnoascus destructans* grows optimally in the cool temperatures at which bats hibernate, with maximal growth at 14°C (20, 21). Bat-to-bat transmission of *P. destructans* is well-established (5, 16), and *P. destructans* can survive in the environment long-term in the absence of bat hosts (22-24). The presence of *P. destructans* in caves and mines is thought to enable seasonal epizootics of WNS, as bats clear infections when they are euthermic during summer (25, 26). However, while it is assumed that exposure to environmental *P. destructans* alone is sufficient to cause WNS in naive bat populations, this remains unproven.

Long-term persistence of *P. destructans* in the hibernacula environment in the absence of bat hosts makes management of WNS challenging as it eliminates the possibility of recolonization of hibernacula with unexposed bats following population extirpation, and reduces the probability that sites will naturally become decontaminated during the summer when bats are no longer inhabiting the site. Additionally, persistence of the pathogen in the environment could facilitate spread to new hibernacula during fall swarm when bats make repeated visits to multiple hibernacula. Here, we assess the role of the hibernaculum as a sufficient reservoir for *P. destructans* to investigate whether transmission of *P. destructans* can occur to naive hosts directly from the environment.

## METHODS

On October 27, 2009, we translocated 79 little brown bats (*Myotis lucifugus*) from a *P. destructans* negative hibernaculum in Wisconsin to two *P. destructans* contaminated mines in Vermont (GM, BWM) from which native bats had been excluded. Collection of live bats was conducted by Wisconsin DNR personnel in compliance with state Endangered and Threatened Species Laws (State Statute 29.04 and Administrative Rule NR 27). In Vermont, handling of bat species was conducted by Fish & Wildlife Department personnel in compliance with Vermont statutes of Chapter 123: Protection Of Endangered Species. New York personnel assisted in live bat handling under the authority of the State of New York State Environmental Conservation Law Article 11.

The source hibernaculum for the *M. lucifugus* used in the study was a mine in northwest Wisconsin, which was 1300 kilometers from the nearest *P. destructans* contaminated hibernaculum at the time of study. GM in Vermont had been confirmed WNS affected in spring of 2008, and BWM was confirmed to harbor bats with WNS in spring of 2009 based on visual inspection of bats and conspicuous mortality. Both sites were straight mining adits, which are small prospecting mines used to explore for mineral deposits, and typically are small with few cracks and crevices and were selected for the simplicity of finding and accessing bats. In July 2009, prior to the experiment, we constructed two bat proof-screens spaced 10 meters apart inside the entrances of both sites. Screens were composed of wooden frames covered in hardware cloth and sealed into the mines using foam sealant and steel wool. After construction of the screens, no native bats were detected in GM during several subsequent visits. At BWM, native bats were able to enter the site up until October 05, 2009 because of a small gap between the ceiling and the first bat proof screen, which allowed access, although bats were not able to pass through the second screen. No native bats were detected in either site after the screen was repaired. At both sites, there were at least one other known hibernacula <1 km from GM and BWM, thus allowing any excluded resident bats to select alternate roosting sites. To ensure recovery of all translocated bats in the experimental portion of the mines, deep crevices (principally at BWM) and drill-holes (principally GM) were plugged or partially filled with roof ridge vent material.

In early October 2009, prior to the introduction of naïve bats from Wisconsin, we collected samples from BWM and GM for microscopic examination and mycological culture. Sterile polyester-tipped swabs were used to sample surfaces where bats were likely to roost (e.g. boreholes) and surfaces that were expected to accumulate *P. destructans* falling from roosting bats or deposited by air currents (e.g. tops of rocks on the mine floor and wall shelves). These sampled sites were located in areas of the mine where concentrations of bats had been observed by state personnel to roost in previous years. Matter collected on the swabs was deposited in 2 ml sterile distilled water in sterile 15 ml centrifuge tubes. Paired swabs of the same targets were then used to streak 100-mm diameter petri plates containing Sabouraud dextrose agar containing gentamycin and chloramphenicol. On return to the laboratory, one drop of solution from the tubes was spread onto a second plate (Sabouraud dextrose agar with gentamycin and chloramphenicol) before the remaining solution was preserved with 1 ml 10% formalin. Media plates were incubated at 5°C.

As a sensitive and specific qPCR was not yet available (e.g. (27)), we used microscopic examination of samples to identify *P. destructans* in accordance with published morphology (18, 20). Prior to microscopic examination, swab solutions were agitated, then centrifuged for 15 minutes at high speed. All but approximately 0.2 ml of the supernatant was carefully discarded with a disposable pipette. The pellet was then resuspended by pipette and, after allowing some of the denser sediment to settle out (1-3 min), 2 drops (0.03 ml) of the fluid was placed on a microscope slide, covered with a 22 × 22 mm coverslip and examined at 450X. The slides were searched systematically by a single observer until at least a single conidium of *P. destructans* was observed. The number of conidia present was then characterized by counting all such conidia on 5 transects across the slide (near top and bottom margins, across the middle, and at the ¼ and ¾ transects).

Many precautions were taken to assure that that the Wisconsin bats were not exposed to *P. destructans* before they were released in the Vermont mines. Naïve bats from Wisconsin were collected by Wisconsin state agency personnel that had never visited any *P. destructans* contaminated sites. All supplies or equipment were either purchased new or disinfected with a 10% chlorine bleach solution. All bats were handled with disposable gloves, one pair per bat. All personnel showered and changed into new clothing before making the trip in a vehicle never before used by anyone who had been to a *P. destructans* contaminated site. Based upon annual sampling of bats, the Wisconsin mine from which the bats originated did not become positive for *P. destructans* until 2016 (7 years after the sampling effort for this experiment was completed), providing strong support that bats were not exposed to *P. destructans* in their origin site at the time they were collected.

Seventy-nine total bats were released into Vermont hibernation sites (n = 38 to BWM, n = 37 to GM). After releasing the bats into the Vermont hibernacula, the sites were checked four times, at intervals of 3, 4, 6, and 8 weeks post introduction (Table 1). At each visit, the hibernacula were systematically searched for live and dead bats. The visual appearance of each bat was noted, as was its exact location. Each bat was also photographed with a high quality digital SLR camera. Except for a careful collection of visible fungus on 3 bats at GM using a polyester swab on the first visit to confirm *P. destructans*, live bats were not physically disturbed. Moribund bats, a status determined by a combination of appearance, location, and reaction to stimuli, were euthanized by cervical dislocation by state agency personnel. Prior to necropsy the bats were weighed and a swab sample was collected from the dorsal surface of the right wing and the entire uropatagium. The swab sample was deposited in 2 ml of distilled water, fixed with the addition of 1 ml 10% formalin, and centrifuged to concentrate conidia and other solids. All but 0.2 ml of the supernatant was then discarded. The pellet and residual fluid were then mixed, and a drop of the mixture placed on a microscope slide and covered with a 22 mm X 22 mm coverslip. Slides were examined systematically for conidia of *P. destructans* at 450X. Once a definitive conidium was detected, a count of conidia was made on three transects as an index of abundance as described above. Histopathological assessment of tissue from the plagiopatagium (18) as well as PCR to confirm presence or absence of *P. destructans* in wing tissue (28) was conducted by the USGS National Wildlife Heath Center. A mean WNS histologic severity score was assigned to each bat for which histopathological assessment was completed (citation 19, appendix S2).

**Table 1.** Progress of white-nose syndrome (WNS) in bats from Wisconsin introduced into bat-free hibernacula in Vermont with histories of WNS outbreaks*

| Location | No. bats seen alive/ No. live bats with visible signs of *P. destructans*†/ No. found dead or moribund |  |  |  |  |
| --- | --- | --- | --- | --- | --- |
|  | December 15 | January 27 | February 18 | March 18 | April 8 |
| BWM | 28/1/8 | 22/17/6 | 16/14/2 | 4/4/16 | 0/0/3 |
| GM | 26/16/4 | 5/4/21 | 0/0/9 | 0/0/3 |  |

\*WNS was first recorded at BWM mine late in the previous winter. WNS was present at GM during the 2 previous winters, after which most bats had died.
†As determined by high-resolution photography.

## RESULTS

### *P. destructans* in Hibernacula before Introduction of Wisconsin Bats

Conidia of *P. destructans* were observed in all 5 samples from drill holes at GM (0.4, 2, 3, 6.6, 67 conidia/transect). Conidia of *P*. destructans were not observed in seven of eight other samples at GM. The single, positive sample from this group was a swab of a rock on the mine-floor sprinkled with bat feces that registered <1 conidium/transect. At BWM, where boreholes are absent, two of 13 samples were positive (0.2 and 0.8 conidia/transect), both from surface swabs of bat carcasses on the mine floor. All culture attempts at both mines were quickly overgrown with other fungi.

### Hibernacula Monitoring

Infection with *P. destructans* was confirmed by photography and microscopic examination of swab samples of bats at both mines by the first visit on December 15, 2009 (Table 1, Fig 1A). Mortality was observed at both mines at this time, although it possible that this mortality was related to or exacerbated by the stress of translocation and not directly caused by *P destructans*. Nonetheless, 16 bats at GM and 1 bat at BWM had visible fungal growth on their skin consistent with *P destructans* infection. Extensive mortality consistent with WNS was recorded at GM in late January 2010 (Fig 1B). No live bats were seen at GM after the February visit. WNS developed significantly more slowly at BWM (Table 1, logistic regression of mortality between sites (GM: -3.342+/-0.584, BWM: -0.538 +/-0.184, P = 0.0021). A single moribund bat at this mine was still alive on the final visit to BWM on April 8, 2010.

**Figure 1.**
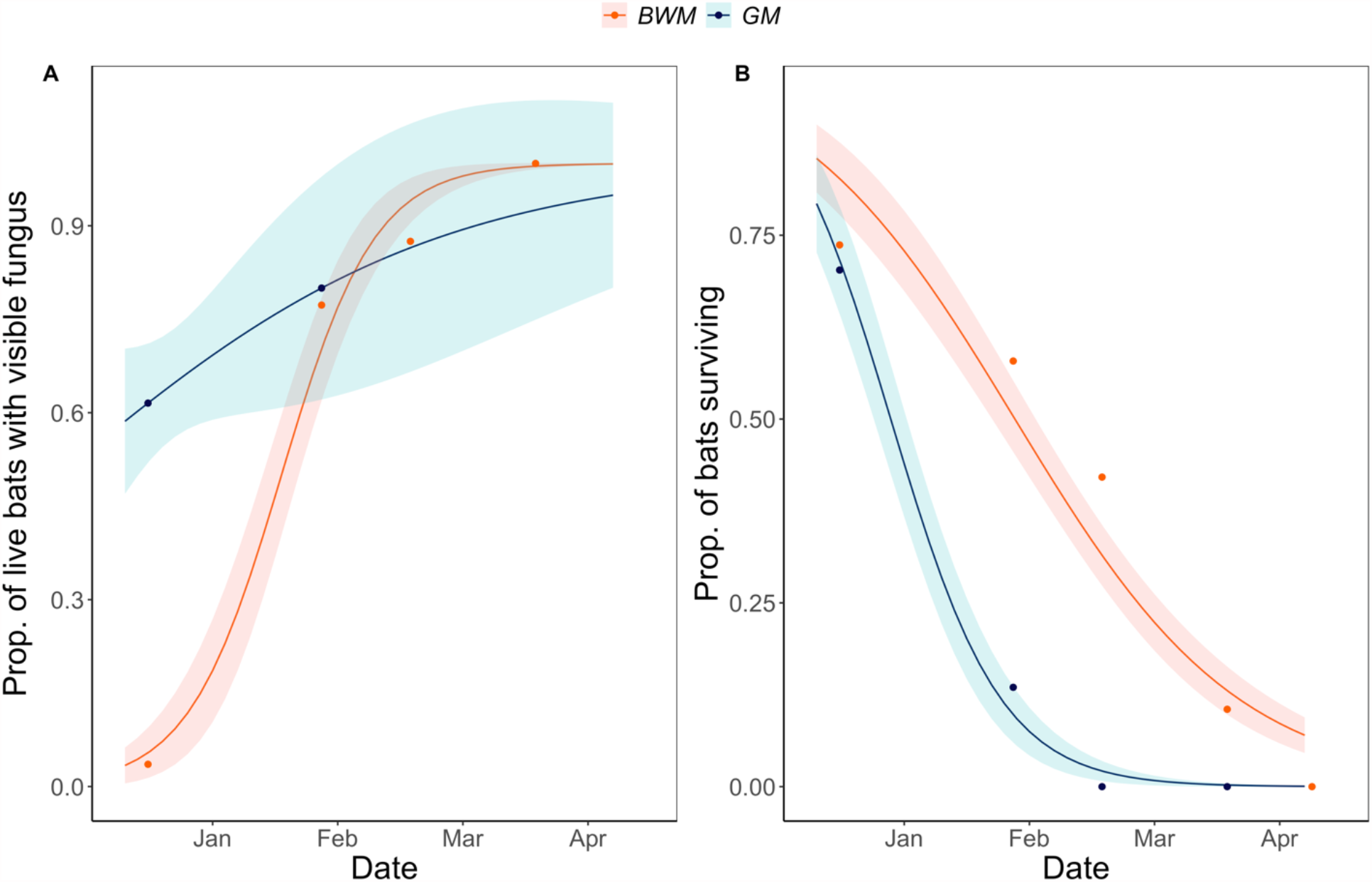
Visible infection and mortality data from the 2009 translocation experiment. (A) The proportion of live bats with visible fungal growth indicative of *P. destructans* infection. (B) The proportion of live bats remaining at each site. Sites differed significantly in their dynamics (logistic regression of site interacting with date, visible fungus site*date coef +/- SE of GM compared to BWM = -2.02 +/- 1.03, P = 0.05, proportion alive at GM compared to BWM: -1.15 +/- 0.41, P = 0.00546).

Most dead bats were recovered toward the front of the mine tunnels (35 bats, 75%, were within 3 m of the screens). Whereas bat that were still alive were encountered in areas where bats previously roosted, regardless of visibly apparent infections with *P. destructans*. Only three non-moribund bats were recorded within 3 m of the screen.

### Confirmation of *P. destructans* and evidence of WNS

Of the 50 carcasses that were suitable for histopathological examination, 45 (90%) showed skin lesions diagnostic of WNS (Figure 2). Five bats lacked diagnostic lesions, 4 of which were recovered on the first visit to BWM, supporting that some initial mortality may have been related to transportation stress. All bats positive for WNS by histopathology were positive for *P. destructans* by microscopic examination of the swab samples for conidia and by PCR of skin samples from the wings. Of the 25 bats with a degree of post-mortem degradation that precluded histopathological assessment, *P. destructans* was detected by swab examination on 17 and by PCR on 18. Subcutaneous white fat was totally or severely (≤0.06 g) depleted in all but 2 of 40 histologically positive bats for which this metric was assessed.

**Figure 2.**
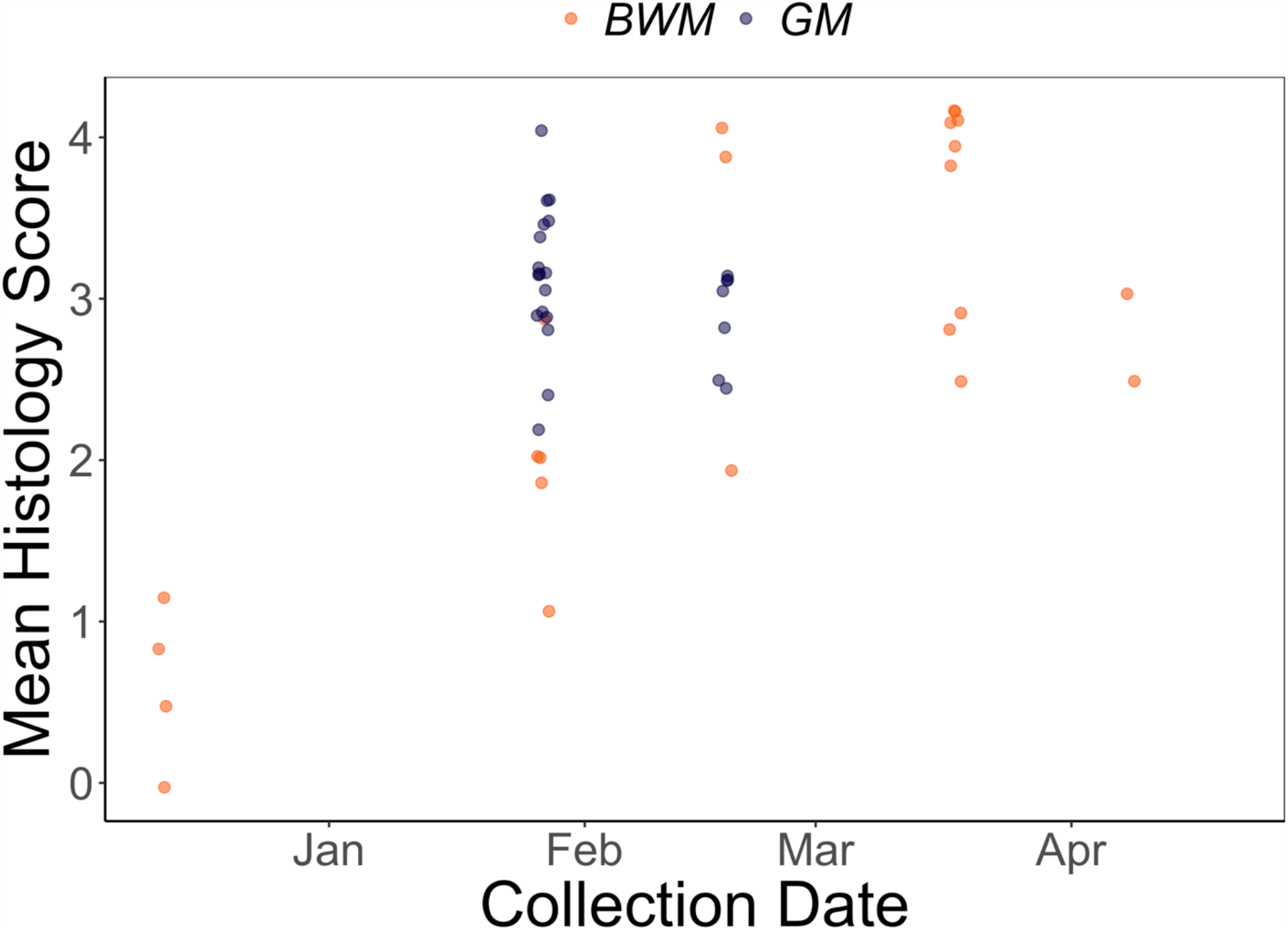
Mean WNS histologic severity scores of dead or moribund bats collected from BWM and GM. Scores are averaged across body surfaces examined (wing, ear/muzzle). Scores were graded as 0 - no fungi suggestive of WNS, 1 - superficial and limited but suspicious of early WNS with hyphae in keratin and randomly into epidermis, but not yet forming distinctive cupping or dense packets, 2 - More extensive superficial infection with epidermal cupping packed with hyphae diagnostic of WNS, 3 - More severe fungal infection with tissue invasion including epidermal cupping packed with hyphae diagnostic of WNS, 4 - Severe infection with tissue and wing damage worse than 3.

## DISCUSSION

Our results indicate that *P. destructans* in WNS-affected hibernacula can serve as a primary source of infection for bats and confirms that the environmental reservoir alone is sufficient to induce infection and mortality with *P. destructans*. The presence of *P. destructans* in a sustained environmental reservoir increases the probability that infection of bats will continue even as bat densities decline, and greatly increases the probability of the complete extirpation at some sites, as has already been documented throughout the eastern U.S. (15, 29, 30). Cumulative losses of hibernating colonies could lead to regional extirpations and increase the potential for species extinction.

Previous work has demonstrated that *P. destructans* contamination in the environment increases with time since *P. destructans* invasion (10, 31, 32) and that infection severity and impacts to host populations increase with the extent of environmental contamination (10). Our findings are similar, in that GM, with a longer history of WNS in bat populations, had a higher number of samples contaminated with *P. destructans* than samples collected from BWM, which is consistent with increasing contamination of hibernation sites over time since *P. destructans* invasion (10, 31, 32). Bats at GM also experienced a faster rate of decline and became visibly infected earlier than bats at BWM, providing additional anecdotal support of the scaling of reservoir contamination and disease impacts. Although this study was limited to only two sites that varied in environmental *P. destructans* contamination and other factors may contribute to differences in impacts (e.g. reviewed in (13)), these data provide support for the potential importance of reservoir contamination in WNS population declines.

Although it is possible that various sources of stress associated with translocating bats in this experiment contributed to the rate of WNS development in our experiment, visible clinical signs of WNS appeared at 49 days post-introduction, earlier than has been documented in laboratory experimental infections, which utilized similar transportation protocols and that may have exerted similarly stressful conditions (16). Many subsequent experimental infections, which confined bats in incubators (e.g. (16, 33, 34), failed to detect such severe clinical signs (e.g. visible fungal infections) as early as was evident in this study. Additional research is needed to determine the underlying differences between experimental and field outcomes.

Critically, our results unequivocally demonstrate that *P. destructans* does not need to be carried by summer bats to cause WNS outbreaks equivalent in scale to those that naturally occur in bat populations. During the summer, prevalence and fungal loads on bats decay (25, 26) and bats become infected upon return to hibernacula during fall (25, 35). While *P. destructans* infections during summer are greatly reduced, viable conidia can be found on small numbers of individuals over summer (36). However, the high infection and mortality in naïve bats in this study demonstrates that recrudescing summer infections are not necessary to initiate epizootics of WNS.

This study was conducted one year after the initial recognition that mass mortality of bat populations in the northeastern U.S. was associated with the fungus *P. destructans* (14). Accordingly, many diagnostic tools and approaches that are now commonly used to assess WNS, such as qPCR to detect the pathogen and UV fluorescence to diagnose fungal lesions, were unavailable to the researchers conducting this work. Subsequent field studies have demonstrated that hibernacula can serve as long-term reservoirs for *P. destructans* (10, 23, 24, 31, 32, 37). However, this study remains the only experiment to assess whether the environmental reservoir can cause WNS epizootics in the absence of previously infected bat hosts. Integrating these experimental data with earlier field studies solidifies the key role of contaminated environments in eliciting WNS outbreaks. More broadly, our results suggest that pairing experiments and field studies can substantially improve understanding of the importance of environmental reservoirs across host-pathogen systems.

## ACKNOWLEDGEMENTS

The authors thank the numerous individuals that contributed to the advancement in knowledge of WNS since this study was first conducted. The use of trade, product, or firm names is for descriptive purposes only and does not imply endorsement by the U.S. government. The findings and conclusions in this article are those of the authors and do not necessarily represent the views of the U.S. Fish and Wildlife Service.

